# pyconsFold: A fast and easy tool for modelling and docking using distance predictions

**DOI:** 10.1101/2021.02.08.430195

**Authors:** J Lamb, A Elofsson

## Abstract

**Motivation:** Contact predictions within a protein has recently become a viable method for accurate prediction of protein structure. Using predicted distance distributions has been shown in many cases to be superior to only using a binary contact annotation. Using predicted inter-protein distances has also been shown to be able to dock some protein dimers.

**Results:** Here we present pyconsFold. Using CNS as its underlying folding mechanism and predicted contact distance it outperforms regular contact prediction based modelling on our dataset of 210 proteins. It performs marginally worse than the state of the art pyRosetta folding pipeline but is on average about 20 times faster per model. More importantly pyconsFold can also be used as a fold-and-dock protocol by using predicted inter-protein contacts to simultaneously fold and dock two protein chains.

**Availability and implementation:** pyconsFold is implemented in Python 3 with a strong focus on using as few dependencies as possible for longevity. It is available both as a pip package in Python 3 and as source code on GitHub and is published under the GPLv3 license.

**Supplemental material:** Install instructions, examples and parameters can be found in the supplemental notes.

**Availability of data:** The data underlying this article together with source code are available on github, at https://github.com/johnlamb/pyconsfold.

## 1 pyconsFold

De novo protein modelling has recently seen significant improvements by relying on contact predictions that have been presented in binary format, two residues are either in contact or not. However, today the best methods are leveraging distance predictions (CASP13 [4, 7]) providing a higher accuracy of the models than if binary contacts were used. To generate a model it is necessary to feed the contact/distance maps into a modelling program. One of the most popular approaches is CONFOLD [1], which is a wrapper around CNS [2, 3] that uses predicted binary contacts together with predicted secondary structure to model proteins. Here, we introduce pyconsFold, a re-implementation and extension of CONFOLD that achieves better results using distance predictions and that also expands to allow for more geometric restraints, such as angles predicted by tools such as trRosetta [7]. Finally, pyconsFold introduces the first easily accessible method for fold-and-dock of two protein chains from inter-chain contacts.

## 2 Modelling

pyconsFold uses predicted distance between pairs of amino acid residues in a sequence. These distances together with either predicted or fixed errors are translated into geometric constraints that is used together with CNS to model the full protein structure. pyconsFold can also be run in contact mode which simulates using binary contact predictions without a predicted distance, this is basically identical to CONFOLD. If side chain angles, for instance predicted by trRosetta, are present, they can also be used as input for further geometric constraint. We have, however, not seen any significant improvement in the model quality using the angles as constraints.

## 3 Docking

pyconsFold introduces a new way of de novo docking together with folding. By using contact predictions which contains both inter- and intra-residue contacts both folding and docking can be done simultaneously. Distance predictions of this type can be done by horizontally concatenating two multiple sequence alignments (MSAs) from two different chains in a complex and adding a poly-G region in between. The poly-G region prevents any spurious false predictions between the end of the first chain and the beginning of the second solely based on proximity. The poly-G region would be trimmed away and residues renumbered before the input is ready for pyconsFold. Using this method, with high quality distance predictions, high quality model and docking can be achieved for some proteins, see Figure 1A). However, the quality of models is strongly dependent on the quality of the MSA and a full study of this is beyond the goals of this paper.

**Figure 1:**
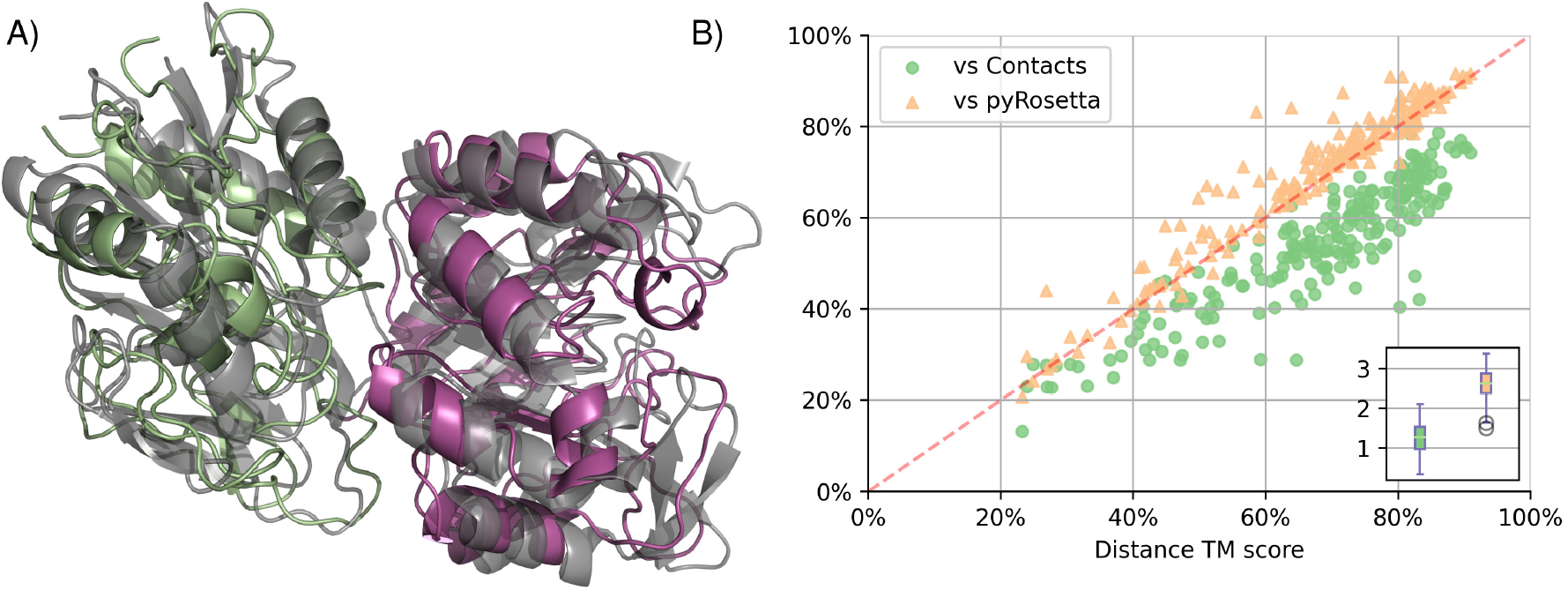
**A**) Model of chain A and B of the 1GPW protein. Using predicted inter- and intra-distances from trRosetta it achieves a TMscore of above 0.8 on each chain with a total RMSD of 3.19A. **B**) pyconsFold distance prediction against both contact (green circles) and pyRosetta (orange triangles) models. Distance predictions outperforms contact predictions in almost all cases. pyconsFolds distance predicted models perform almost as well as pyRosetta but is around 20 times (more than 1 order of magnitude) faster per model as the inset shows(loglO scale of per model time in seconds. pyconsFold in green and pyRosetta in orange.

## 4 Additional features

For ease of use and reproducibility we have included several extra features and utilities. By default the generated models are ranked by CNS internal NOE energy, but Quality Assessment score *pcons* [5] will also be calculated. If a native structure is known and supplied with the tmscore_pdb_file argument, the *tmscore* [8, 6] for each model against the native structure will be calculated. Compiled versions of both pcons and TMscore for unix based x64 systems are packaged together with pyconsFold under the open source Boost license. If your system does not support the built in versions, you can manually install them on your system and as long as they are in your path, will be chosen instead of the built in binaries.

## 5 Conclusion

pyconsFold offers a complete toolkit for de novo modelling using predicted contact distances and angles. Its focus is on ease of use and reproducibility and is available both as source on github and as an easily installable pip package in Python 3. It comes packaged with QA-programs to rank the generated models and allows transparency of parameters for the underlying CNS-system. It also offers an innovative de novo fold-and-dock protocol where predicted inter chain contacts are used as restraints for docking. It offers a significant increase in accuracy over contact based protocols by using predicted distances, Figure 1B), and a comparable performance to pyRosetta although being around 20 times faster.

## Supporting information

Supplemental installation instructions

## Acknowledgements

This work was supported by grants from the Swedish Research Council (VR-NT 2016-03798) and SNIC to A.E.

