## Supplemental installation instructions for "pyconsFold: A fast and easy tool for modelling and docking using distance predictions"

### pyconsFold: Supplemental Notes

Lamb J., Elofsson A.

#### 1 Installation

pyconsFold requires a working installation of CNS. This needs to be done manually due to license.

##### 1.1 CNS

1. Request a download link from CNS.
2. Follow the emailed instructions to download
   cns_solve_1.3_all_intel-mac_linux.tar.gz
3. Extract the files
   tar xzvf cns_solve_1.3_all_intel-mac_linux.tar.gz
4. Change into the resulting directory
   **cd** cns_solve_1.3
5. Unhide the bash-specific file
   mv .cns_solve_env_sh cns_solve_env.sh
6. In this resulting file, replace _CNSsolve_location_ with the CNS installation folder. If you extracted the file in your home folder then the CNS installation would be:
   /home/<your username>/cns_solve_1.3
7. Source CNS, source cns_solve_env.sh[1], to make this permanent and to prevent you having to do this every time, add it to your .bashrc file.
8. Test CNS by going into the test folder cd test and run the tests
   ../bin/run_tests -tidy *.inp

##### 1.2 pyconsFold

1. Run:
   pip3 install pyconsFold
   It is recommended to run a separate python environment for pyconsFold, ie virtualenv.

[1] If you get an error about csh interpreter, you need to install csh

### 2 Examples

#### 2.1 Distance modelling

Distance modelling is done using a contacts file with predicted distances and errors.

```
import pyconsFold
pyconsFold.model_dist(fasta_file, contact_file,
                      output_directory)
```

#### 2.2 Classical modelling

Classical model is done by treating the contacts file as binary contacts and using a static distance and error. This means a contacts file with predicted distances can be used as input but the program will treat the contacts as static contacts with the same distance and error.

```
import pyconsFold
pyconsFold.model(fasta_file, contact_file,
                 output_directory)
```

#### 2.3 Docking modelling

Docking requires a contacts file with both inter and intra contacts between the two proteins. If no inter contacts are found, a warning is generated and an artificial contact is created between the centers of the two proteins to prevent the program from failing. Either predicted distances or classical contacts can be used, see the dist parameter.

```
import pyconsFold
pyconsFold.model_dock(first_fasta_file,
                      second_fasta_file,
                      contact_file, output_directory)
```

### 3 Arguments

Multiple optional arguments are available. The following are the most commonly used. For a full list, see the github repository.

```
rr_pthres  -- Threshold for the confidence we
              want in a prediction (default
              model(0.80), model_dist(0.45),
              model_dock(0.50))
rr_sep     -- Separation between contacts
              (default 0)
```

```
save_step  -- Save working steps
              (default False)
stage2     -- Run stage2, filter contacts vs
              generated structure and generate
              new structures with filtered
              contacts (default False)
debug      -- Write out debug information
              (default False)
selectrr   -- How many contacts to use?
              Can be "all", "#L", or #.
              (default "all")
mcount     -- How many models to generate?
              (default 20)
top_models -- How many of the generated
              models should be ranked and
              saved? (default 20)
use_angles -- If predicted angels should be
              used, only works with npz
              (default False)
omega      -- RR-formated file with omega
              angles (if npz are not used)
              (default '')
theta      -- RR-formated file with theta
              angles (if npz are not used)
              (default '')
```

### 4 Arguments

QA-function arguments to all above functions:

- pcons (default False) – If set to true, gives pcons scores for all models (using either pcons installed in the PATH or the builtin binary)
- tmscore_pdb_file – If a structure file is supplied, runs all models against this (presumed) native structure and reports the TMscore (using either TMscore in the PATH or builtin binary)

### 5 Extras

```python
from pyconsFold.utils import npz_to_casp, pdb_to_npz

npz_to_casp("trRosetta.npz")  ## Converts trRosetta
                              ## distance and angle
                              ## predictions to CASP
                              ## format in separate files
```

```
pdb_to_npz("structure.pdb")    ## Converts a structure
                               ## (pdb/mmCif) to trRosetta
                               ## distances and angles,
                               ## useful when
                               ## investigating how well
                               ## a model conforms to
                               ## restraints
```

### 6 Adjustable parameters for CNS, advanced

```
rrtype      -- Between which atoms in a residue are the contacts?
               (default 'cb')
lbd         -- Lambda, 0.1-10 (default 0.4)
contwt      -- Contact restraint weights, 0.1-10000 (default 10)
sswt        -- Secondary structure weights, 0.1-100 (default 5)
bin_values  -- Dictionary of bin_values for conversion of npz to
               RR-format, see source code (default {})
```
